# Genotype by microbiome interactions have large effects on growth in *Lotus japonicus*

**DOI:** 10.1101/2022.03.28.486086

**Authors:** Masaru Bamba, Turgut Yigit Akyol, Yusuke Azuma, Johan Quilbe, Stig Uggerhøj Andersen, Shusei Sato

## Abstract

Biological interactions between plants and their root microbiomes are pivotal for plant growth. Even though the plant genotype [G], soil microbiome [C], and growth conditions (environment) [E] are core factors shaping the root microbiome, their relationships remain unclear. We disentangled the effects of G, C, E, and their interactions on the *Lotus* root microbiome and plant growth using a cross-inoculation approach that reconstructed the interactions between nine *Lotus* accessions and four soil microbiomes under two different environmental conditions. We found that a large proportion of the root microbiome composition was determined by C and E and that G-related (G, G × C, and G × E) effects were significant but small. In contrast, the interactions between G and C had a more pronounced effect on plant shoot growth than C alone. Our findings indicate that most microbiome variations controlled by C have little effect on the plant phenotype, whereas G × C interactions have more significant effects. Plant genotype-dependent interactions with soil microbes warrant more attention in efforts to optimize crop yield and resilience.

## Introduction

The interaction between the microbiome and plant roots is one of the most influential factors affecting plant growth. This interaction is pervasive and can have an extensive effect on host plants, such as disease resistance (Santhanam et al., 2015; Busby et al., 2016; Carrion et al., 2019), stress tolerance (de Vries et al., 2019; Liu et al., 2020), nutrient supply (Zhang et al., 2019), and overall plant health (Berendsen et al., 2012). Consequently, there has been increasing interest in clarifying how plant-microbiome interactions are established, maintained, and exploited in agronomy and ecology (Mauchline and Malone, 2017). The root microbiome structure results from complex interactions among host plants, soil microbiome, and abiotic environments. A plant recruits bacteria from the soil microbiome to its root/rhizosphere and establishes its root microbiome, which deviates considerably from the soil microbiome in a particular environment; consequently, the root microbiome responds to changes in plant status. For this reason, there is a need to disentangle the interactions among the effects of plants, soil microbiome, and environment to understand the dynamics and function of the root microbiome.

Plant genetic differentiation is one of the most studied plant factors that affect root microbiome structure (Bamba et al., 2019). *Arabidopsis thaliana* host genotypes have a small but significant influence on the microbes inhabiting the endophyte compartment of their roots (Bulgarelli et al., 2012; Lundberg et al., 2012). Similar patterns have been observed in *Medicago truncatula* (Brown et al., 2020), tomato (Weinert et al., 2011), and inbred maize lines (Peiffer et al., 2013; Walters et al., 2018). In the interspecies-level comparisons, the phylogenetic distance between plants and root microbiome dissimilarity appeared to be correlated in Brassicaceae, Poaceae (Bouffaud et al., 2014; Schlaeppi et al., 2014; Terrazas et al., 2020) and higher taxonomic levels (Wang and Sugiyama, 2020), supporting the effects of host genetics on the root microbiome. In contrast, the host genotypes of *Boechera stricta* in field experiments did not show statistically significant effects on their root microbiome structures (Wagner et al., 2016) because of low genetic divergence caused for thousands of years (Rushworth et al., 2011). According to these studies, plant genetic differentiation could drive the divergent host genotype effects on the root microbiome.

The response of plants to different environments also alters their root microbiome (Bouskill et al., 2013), and its impact depends on the plant genotype. Lundberg et al. (2012) concluded that the plant genotype was less important for the root microbiome structures than the soil type containing differences in the microbiome and environment. In contrast, their genotype-dependent effects were also observed (Lundberg et al., 2012). A recent pot experiment showed that the effect of environmental treatment on the root microbiome (both fungal and bacterial communities) varied among plant genotypes (Gallart et al., 2018). Environmental treatments can also alter the microbiomes in the soil, rhizosphere, and root endophytes (Naylor et al., 2017; Yeoh et al., 2016). Accordingly, plant genotype effects on the root microbiome could exist in a complex entanglement with the environment and soil microbiome. However, it remains unclear how the plant genotype [G], soil microbiome [C], and soil environment [E] relate to each other in shaping the plant root microbiome and its impact on plant phenotypes.

Here, to disentangle the effects of G, C, E, and their interactions on plant root microbiome and plant phenotypic variation, we reconstructed their interactions using nine *Lotus* accessions and four soil microbiomes under two different environmental conditions. *Lotus japonicus* is a model species for understanding plant-microbe interactions (Handberg and Stougaard, 1992; Kawaguchi, 2000; Bamba et al., 2019, 2020). The Japanese population has originated and experienced recent population expansion in the Japanese archipelago during the last approximately 20 thousand years (Shah et al., 2020). We chose eight different accessions and one closely related species, *Lotus burttii*, as host plants based on their population genomic information. For the soil microbiome, we focused on two bacterial communities extracted from the soils of the Kashimadai field at Tohoku University, Japan. Each of them alone, a 1:1 mixture and a non-inoculant control, were used in this study. The soils were obtained from two adjacent plots (F5C and F5S) and irrigated using underground water containing ∼1/4 the salt concentration of seawater to F5S and regular water to F5C from 2017 to 2019. These inoculation experiments were performed in environments with and without salt, corresponding to the environment in which the soil microbial community was sampled.

In the present study, we performed 16S rRNA amplicon sequencing using MAUI-seq technologies (Fields et al., 2020) and conducted community analyses. We aimed to quantify the effects of G, C, and E on root microbiomes and identify which microbes are sensitive to plant genotypes. Second, we compared plant terminal phenotypes in the cross-inoculation experiments to quantify these effects on plant growth.

## Results

We performed a cross-inoculation experiment using nine *Lotus* accessions [G] and four inoculants [C] under two conditions [E], resulting in 72 combinations. We collected 768 plant individuals (6–12 per combination) (Supplemental Table S1). Although we cultivated 12 plants for each combination, approximately 9% of plants did not survive.

### Plant root microbiomes and effects of G, C, and E

We investigated the root microbiomes of 327 plants. Using MiSeq sequencing, we obtained 168,097,504 reads (ranging from 100,220 to 830,796 per individual). After pre-processing, 115,530,212 reads were allocated to 327 individuals, ranging from 55,512 to 795,588. We used all quality-filtered reads for counting Unique Molecular Identifiers (UMI). In addition, 38,813 unique sequences, with an average number of UMIs greater than or equal to 0.1, were used for the BLAST search. As a result of the BLAST search, 35,370 sequences consisting of 16,785,973 reads were derived from the bacterial 16S rRNA genes. Bacterial sequences were assigned to 4,225 different bacterial strains, 230 genera, and 70 families (Figure 1).

**Figure 1.**
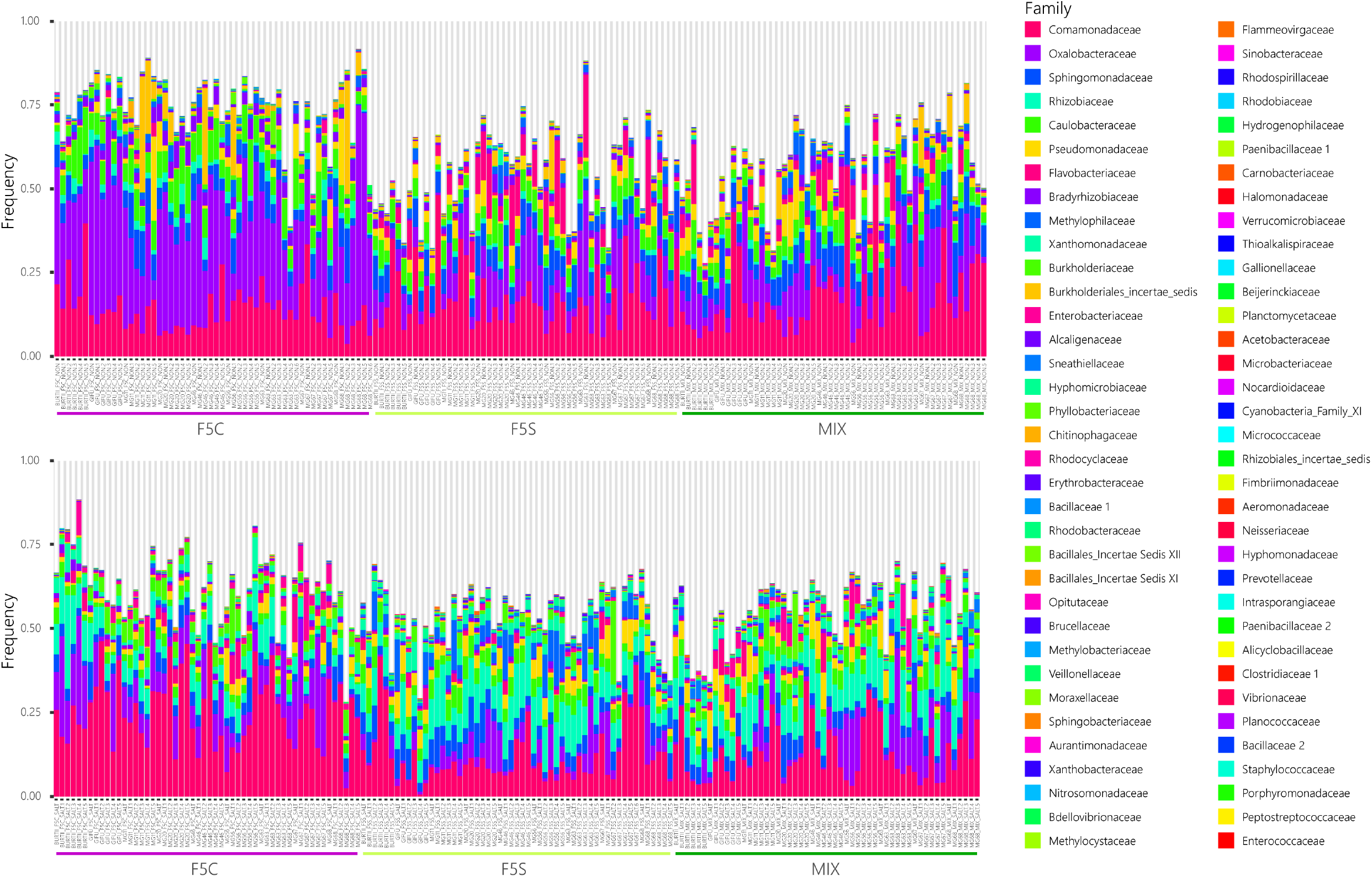
Family level composition of *Lotus* root microbiomes. A color-coded bar plot shows the bacterial family abundance in the *Lotus* root sample. Sample names were given “Genotype”_”Inoculant”_” Environment”_replicate in this study. The upper and lower parts are shown with and without salt, respectively. The grey portion of plots indicated the “NotAssigned” taxa. Arabic and Greek numerals following the family name are based on the classification in RDP11.

Prior to diversity analysis, we performed coverage-based rarefaction to remove bias caused by the different numbers of sequenced reads among samples using the aggregated data based on the BLAST top hit strain. Because the lowest slope at the end of the rarefaction curve among the samples was 0.0270, we resampled all samples so that their slope at the end of the rarefaction curve was equal to that value. (Supplemental Figure S1).

We calculated the α-diversities of the *Lotus* root microbiome, using the Shannon index based on the rarefied composition data, and they ranged from 2.789 to 7.230 (Figure 2). We detected a significant effect of G and C on the α-diversity while their interaction and the environment had no effect (P < 0.05, Supplemental Table S2). The Tukey-Kramer test indicated that the root microbiomes of MG20 were more diverse than those of Gifu and MG68, and the microbiomes with MIX were more diverse than those with F5C inoculants (P < 0.05, Supplemental Table S3).

**Figure 2.**
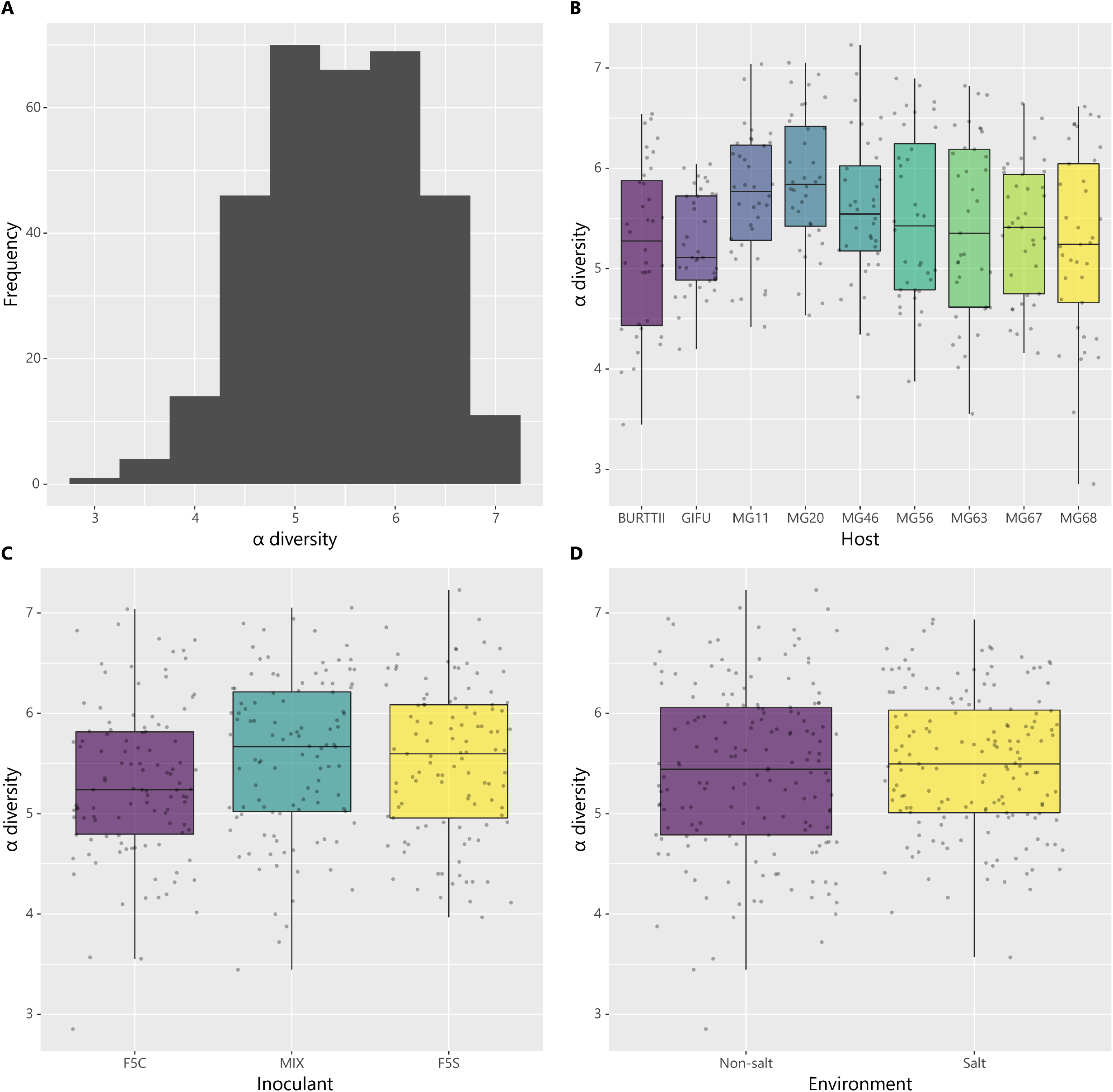
α-diversities of root microbiomes. α-diversity based on the Shannon index with a strain-level taxonomic assignment. (A) Distribution of α-diversity in the *Lotus* root samples. (B, C, and D) Comparison of α-diversity among groups of host genotypes [G], inoculant communities [C], and environments [E], respectively.

The community structures of the root microbiomes were characterized based on the- β diversity (Morisita-Horn index). Non-metric multidimensional scaling (nMDS) analysis showed an apparent difference between environments [E] and among inoculants [C], whereas the differences between hosts [G] were unclear (Figure 3). The PERMANOVA analysis indicated that G, C, E, and their interactions significantly affected the root microbiome structure (P < 0.05, Table 1). The effects of C and E were the largest, explaining about 22% of the variance. The others (G, G × C, G × E, C × E, and G × C × E) were around 4%, and 35% of the variance was residual. This result was comparable to the community structure of the root microbial community within *L. japonicus* species (Supplemental Table S4), indicating that the root microbiome of *L. burttii* accession did not deviate from that of *L. japonicus*. Evaluating the G effect in conditions where the combination of C and E was fixed showed that the differences in G could explain approximately 25-40% of the variation in microbiome composition (Supplemental Table S5). In addition, the differences in the root microbiome among host plants were not correlated with the genetic distances between host accessions (P > 0.05, Supplemental Figure S2; Supplemental Table S6).

**Figure 3.**
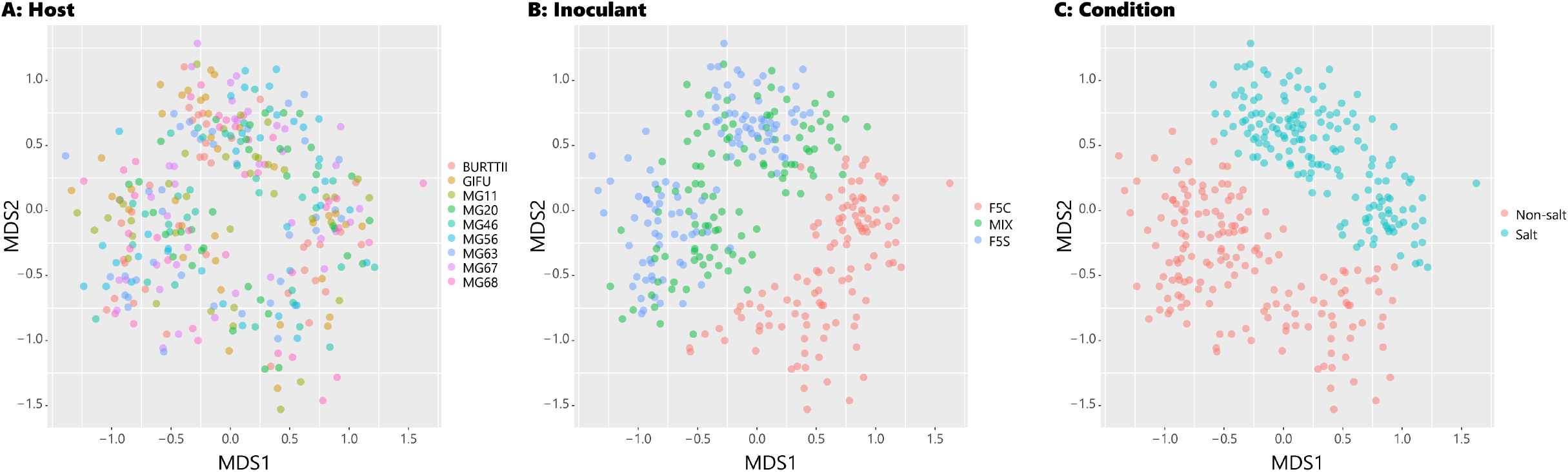
Root microbiome structures based on β-diversity. Non-metric multidimensional scaling (nMDS) for *Lotus* root microbiome dissimilarity (Morisita-Horn index) is shown. (A, B, and C) Colors represent different plant genotypes, inoculants, and conditions.

**Table 1.** Permanova results for variation in root microbiomes.

|  | DF<sup>a</sup> | SS<sup>b</sup> | MSS<sup>c</sup> | F value<sup>d</sup> | R<sup>2</sup> | P value |  |
| --- | --- | --- | --- | --- | --- | --- | --- |
| G | 8 | 4.0915 | 0.5114 | 4.2391 | 0.0436 | 1.00E-04 | *** |
| C | 2 | 20.6411 | 10.3206 | 85.5426 | 0.2199 | 1.00E-04 | *** |
| E | 1 | 20.5804 | 20.5804 | 170.5821 | 0.2193 | 1.00E-04 | *** |
| G × C | 16 | 3.6375 | 0.2273 | 1.8844 | 0.0388 | 1.00E-04 | *** |
| G × E | 8 | 3.9657 | 0.4957 | 4.1088 | 0.0423 | 1.00E-04 | *** |
| C × E | 2 | 3.9470 | 1.9735 | 16.3576 | 0.0421 | 1.00E-04 | *** |
| G × C × E | 16 | 4.0601 | 0.2538 | 2.1033 | 0.0433 | 1.00E-04 | *** |
| Residuals | 273 | 32.9369 | 0.1206 |  | 0.3509 |  |  |
| Total | 326 | 93.8604 |  |  | 1 |  |  |

<sup>a</sup> Degree of freedom. <sup>b</sup> Sums of squares. <sup>c</sup> Mean sums of squares. <sup>d</sup> Pseudo-F value in permutation

To identify which bacterial strains were affected by G, C, and E, we evaluated these effects using a generalized linear model (GLM), in which the response variable was each bacterial frequency. Of the 3,700 strains, 3,333 were significantly affected by any G, C, and E variable and their interactions. The G variable had a significant effect on 1221 of these strains; however, 2485 and 2634 strains were affected by C and E, respectively (Supplemental Table S7; Supplemental Figure S3). The strains affected by G, C, and E were shared by G versus C (33%), G versus E (32%), and C versus E (57%) (Supplemental Figure S4A). The variables containing G (G, G × C, G × E, and G × C ×E) had significant effects on 1,928 strains, and the strains were shared by G vs. G × C (38%), G vs. G × E (52%), and G vs. G × C × E (34%) (Supplemental Figure S4B). Moreover, the enriched genera that were significantly affected by the variables containing G were *Pseudomonas*, *Sphingobium*, *Ralstonia*, and *Delftia* (Fisher’s exact test FDR-P < 0.05, Supplemental Table S8). Similarly, the enriched families affected by the same variables were Enterobacteriaceae, Sphingomonadaceae, Pseudomonadaceae, Burkholderiaceae, and Methylophilaceae (Fisher’s exact test FDR-P < 0.05, Supplemental Table S8).

### Plant phenotype and effects of G, C, E

We obtained four phenotypes (shoot length, SL; root length, RL; number of leaves, NOL; and number of branches, NOB) from all 749 individuals (Figure 4). All phenotypic traits were positively correlated (Pearson’s product-moment correlation: P < 0.001, Supplemental Figure S5). All combinations of phenotypic traits, except for those between RL and NOB, were significantly correlated in the G, C, and E groups (Pearson’s product-moment correlation: P < 0.05, Supplemental Figures S5, S6, and S7). In the cross-inoculation experiment, we detected significant effects of G, C, and E and their interactions on all four phenotypes with GLM, except for C × E on SL and NOB (F test P < 0.05, Supplemental Table S9; Supplemental Figure S8). The most prominent effects on all phenotypes were G × C × E. By contrast, the other effect sizes of each variable in the GLM varied by phenotype: the G effect was the largest on SL and RL, and the E effect was the largest on NOL and NOB (Supplemental Table S9). The coefficients of salt addition, as an E factor, were positive for plant shoot phenotypes, and the Tukey-Kramer test indicated significant differences between salt and non-salt conditions (P < 0.001, Supplemental Table S10) for all phenotypes; that is, NaCl in the growth media promoted plant growth. The C coefficients for F5C, MIX, and F5S in the GLM were mostly negative, except for F5C in RL. The Tukey-Kramer test showed significant differences between non-inoculant and inoculant conditions (P < 0.001, Supplemental Table S10). This result indicates that the microbiomes in the Tohoku fields had adverse effects on plant growth.

**Figure 4.**
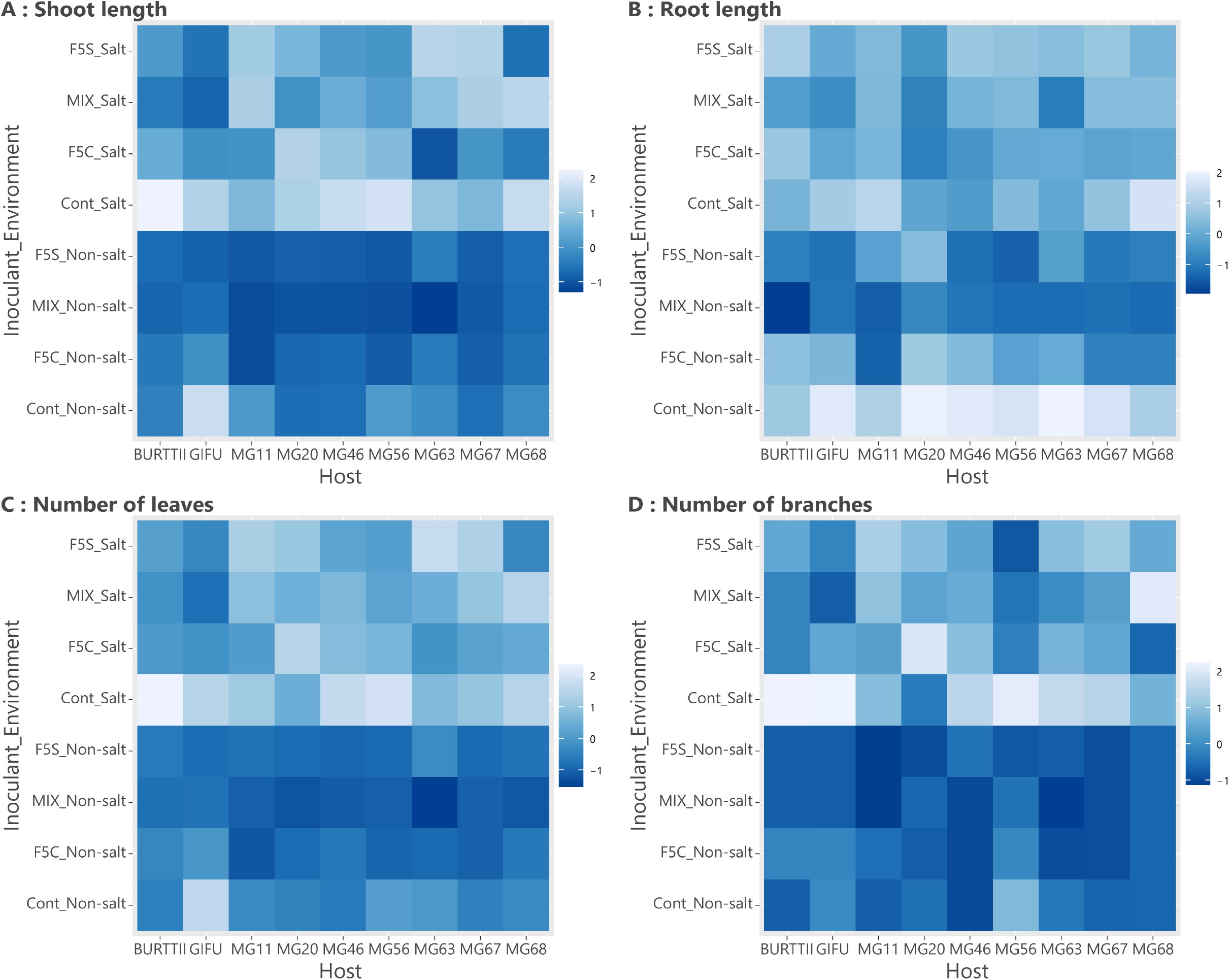
Plant phenotypic variation in the cross-inoculation experiments. Heatmaps of the four plant phenotypes: (A) shoot length, (B) root length, (C) number of leaves, and (D) number of branches. Each cell color indicates standardized phenotypic values for each plant genotype.

The GLM without non-inoculant data showed that all G, C, and E cases and their interactions significantly affected all four plant phenotypes, except for C × E on RL (Table 2; Supplemental Figure S9). The largest effect size for all the plant phenotypes in the model was G × C × E. The second-largest effect on SL, RL, and NOL was on G, while that on NOB was on E. The C variables showed that the differences in inoculant communities in this model had a less significant effect on plant phenotypes (Figure 5). The η^2^ of variable C ranged from 0.003 to 0.03, which is less than or equal to “small” by Cohen 1998’s guideline. The η^2^ of variable G × C was between 0.025 and 0.031, which is assigned to “small”. These η^2^ values of G × C variables were larger than those of C variables for all phenotypes, indicating that G × C variables were more significant than C.

**Figure 5.**
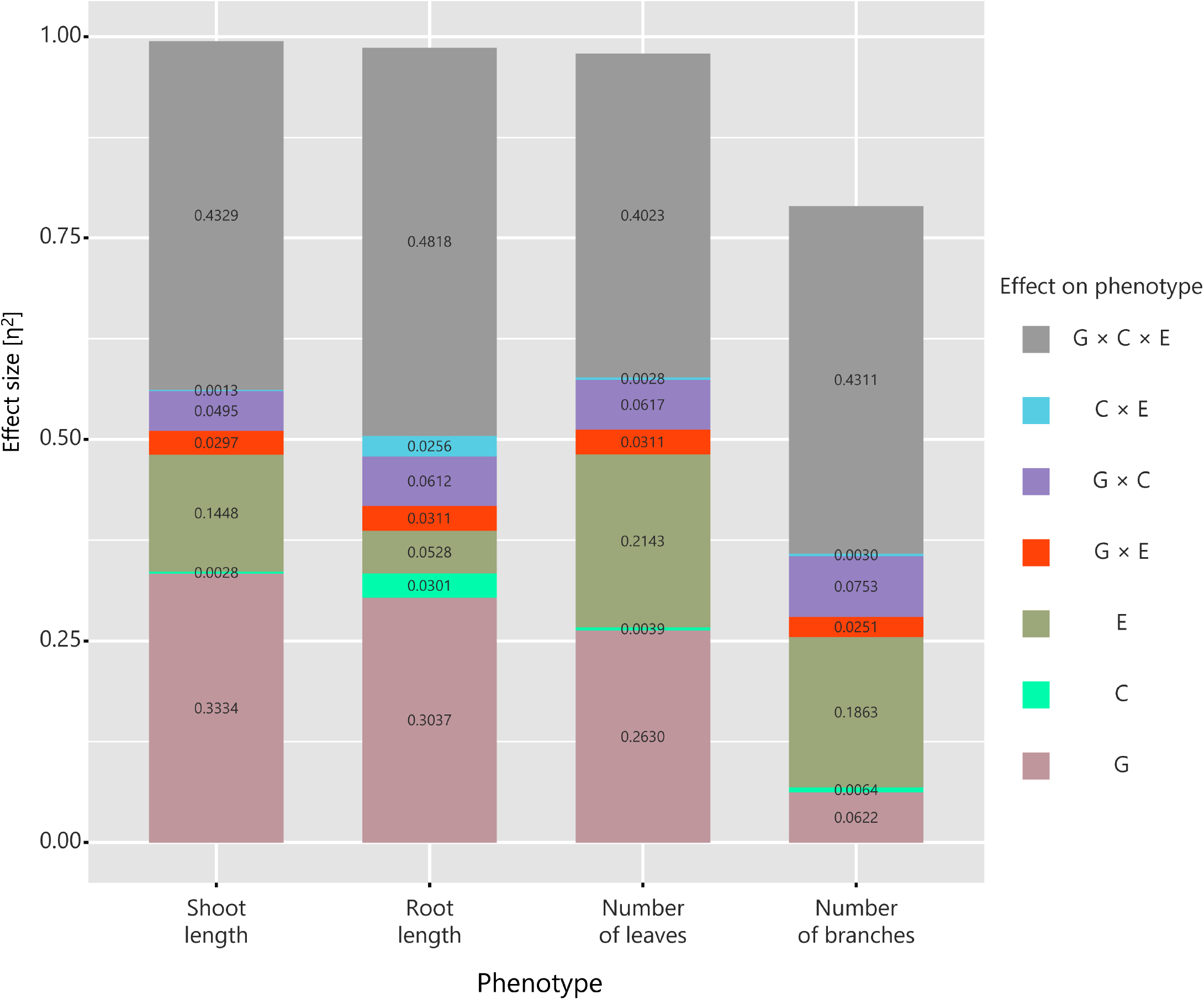
Effect sizes of G, C, E, and their interactions on plant phenotypes. The portion of each color and number on the bar chart represent the effect size η^2^ of each variable in the generalized linear model without non-inoculant data.

**Table 2.** Generalized linear model for plant phenotypes in the cross-inoculation experiment without non-inoculant data.

| | | DF<sup>a</sup> | SS<sup>b</sup> | F value | $\eta^2$ | P value | |
| --- | --- | --- | --- | --- | --- | --- | --- |
| SL<sup>c</sup> | G | 8 | 142.7981 | 89.5157 | 0.3334 | 3.52E-92 | *** |
|  | C | 2 | 2.6835 | 6.7287 | 0.0028 | 1.30E-03 | ** |
|  | E | 1 | 39.6516 | 198.8508 | 0.1448 | 2.24E-38 | *** |
|  | G × C | 16 | 13.0064 | 4.0767 | 0.0297 | 2.21E-07 | *** |
|  | G × E | 8 | 9.6809 | 6.0686 | 0.0495 | 1.79E-07 | *** |
|  | C × E | 2 | 0.6253 | 1.5678 | 0.0013 | 2.10E-01 |  |
|  | G × C × E | 16 | 6.3997 | 2.0059 | 0.4329 | 1.14E-02 | * |
|  | Residuals | 511 | 101.8952 |  |  |  |  |
| RL<sup>d</sup> | G | 8 | 55.8287 | 45.0103 | 0.3037 | 1.34E-54 | *** |
|  | C | 2 | 7.7638 | 25.0375 | 0.0301 | 4.23E-11 | *** |
|  | E | 1 | 13.7103 | 88.4282 | 0.0528 | 1.77E-19 | *** |
|  | G × C | 16 | 12.4633 | 5.0241 | 0.0311 | 9.96E-10 | *** |
|  | G × E | 8 | 9.0006 | 7.2565 | 0.0612 | 3.85E-09 | *** |
|  | C × E | 2 | 5.3485 | 17.2484 | 0.0256 | 5.64E-08 | *** |
|  | G × C × E | 16 | 7.8354 | 3.1586 | 0.4818 | 3.50E-05 | *** |
|  | Residuals | 511 | 79.2275 |  |  |  |  |
| NOL<sup>e</sup> | G | 8 | 66.5305 | 64.1591 | 0.2630 | 2.51E-72 | *** |
|  | C | 2 | 2.4078 | 9.2878 | 0.0039 | 1.09E-04 | *** |
|  | E | 1 | 34.9319 | 269.4944 | 0.2143 | 6.00E-49 | *** |
|  | G × C | 16 | 10.0314 | 4.8369 | 0.0311 | 2.92E-09 | *** |
|  | G × E | 8 | 5.5026 | 5.3065 | 0.0617 | 2.07E-06 | *** |
|  | C × E | 2 | 0.5684 | 2.1927 | 0.0028 | 1.13E-01 |  |
|  | G × C × E | 16 | 4.3638 | 2.1041 | 0.4023 | 7.29E-03 | ** |
|  | Residuals | 511 | 66.2358 |  |  |  |  |
| NOB<sup>f</sup> | G | 8 | 5.5351 | 5.7189 | 0.0622 | 5.51E-07 | *** |
|  | C | 2 | 1.1457 | 4.7350 | 0.0064 | 9.17E-03 | ** |
|  | E | 1 | 22.9586 | 189.7658 | 0.1863 | 6.12E-37 | *** |
|  | G × C | 16 | 3.5567 | 1.8374 | 0.0251 | 2.41E-02 | * |
|  | G × E | 8 | 7.3823 | 7.6274 | 0.0753 | 1.16E-09 | *** |
|  | C × E | 2 | 0.3147 | 1.3005 | 0.0030 | 2.73E-01 |  |
|  | G × C × E | 16 | 3.4697 | 1.7925 | 0.4311 | 2.92E-02 | * |
|  | Residuals | 511 | 61.8228 |  |  |  |  |

<sup>a</sup> Degree of freedom. <sup>b</sup> Sums of squares.
<sup>c</sup> Shoot length. <sup>d</sup> Root length. <sup>e</sup> Number of leaves. <sup>f</sup> Number of branches.

In this study, the potential confounding factors derived from each pot could have caused an overestimation of G × C × E effects because the pot differences masked all G × C × E combinations. First, we calculated how much variation in plant phenotypes was explained by the differences in pots for each G × C × E combination. On average, 15% and 8% of the variance in SL and RL, respectively, were derived between pot replicates, indicating that variation existed between them. To consider this variance in the analysis, we randomly selected one of the pots from each combination and evaluated the effects of G, C, E, and their interactions on plant phenotypes. Although we could not distinguish the G × C × E effects from pot effects during the permutations, the other effects could be estimated by considering the variability derived from pot effects. The statistical significance represented by P values of G, C, E, G × C, and G × E effects were mostly distributed below 0.05; on the other hand, the P values of C × E effects on plant phenotypes, except for RL, were skewed distributed above 0.05 (Supplemental Figure S10).

Moreover, for SL, RL, and NOL, the effects of the G variable were the largest and those of the E variable were the largest for NOB. The differences in the inoculant communities had smaller effects on all phenotypes than the interactions between the G and C variables (Supplemental Figure S11). Because these results were comparable to the results of GLM with a complete dataset, the estimation of the effects of G, C, and E, and their interactions are likely to be meaningful, even if pot effects are present.

Since both plant phenotypes and microbial communities depend on the effects of G, C, E, and their interactions, we attempted to integrate the variation in SL and root microbiome structure with variance component analysis. We used standardized SL values by the G factor to calculate SL variation because this factor explained a large amount of SL and little root microbiome structure. Variation in the root microbiome structure was calculated based on 1 - the Morisita-Horn similarity index matrix, an identical matrix used in the community analysis. In this analysis, 55% of the variance in SL could be explained by the similarity of root microbiome structures. This result indicated that identifying which microbes could affect plant growth was difficult, even though many kinds of microbes in the soil microbiome would have favorable or adverse effects on plant phenotypes.

## Discussion

While there have been several attempts to evaluate the individual effects of plant genotype [G], inoculant community [C], and growth condition [E], studies comparing these effects and their interactions on the plant root microbiome and phenotypes have been uncommon. Here, we performed cross-inoculation experiments using nine *Lotus* accessions and four inoculant microbial communities under two conditions and characterized plant phenotypes and root microbiomes. The cross-inoculation experiments were conducted in controlled environments and enabled us to disentangle the effects of G, C, and E, and their interactions on the *Lotus* root microbiome and phenotypes.

In the cross-inoculation experiments, the microbiome detected in plant roots had the features of a root/rhizosphere microbial community. The largest proportion of the microbial community was Proteobacteria, followed by Bacteroidetes and Firmicutes, with these three Phylums accounting for approximately 90% of the community (Figure 1; Supplemental Table S11). Proteobacteria is one of the most enriched phyla in the plant rhizosphere compared with the soil microbiome (Peiffer et al., 2013). Firmicutes and Bacteroidetes are the predominant phyla in the root microbiome (Guo et al., 2017; Enebe and Babaloa, 2020). Consequently, we identified the microbiome that inhabited the *Lotus* root and rhizosphere in this study.

Meanwhile, our analysis detected few Actinobacteria commonly observed in the rhizosphere (Yadav et al., 2018). Actinobacteria were observed in the plant roots during our preliminary experiment, in which *Lotus* was grown directly at the site where we collected the inoculated community used in this study (Bamba unpublished; data not shown). The small number of Actinobacteria may be explained by our inoculation methods. The growth pots were filled with vermiculite and media and kept anaerobic, and these conditions are unfavorable for most Actinobacteria that are aerobic (Trujillo, 2016). Therefore, we should note that our cross-inoculation experiment did not reflect the complete relationship between the plant and soil bacterial communities.

The microbial communities used in this study reflected the unique features of our experimental field (F5C and F5S in the Kashimadai field). First, we found that all the microbial communities used in this study had adverse effects on *Lotus* growth (Supplemental Table S10). This result suggests that the soil microbial communities in the Kashimadai fields had enriched pathogenic microbes during three years of *Lotus japonicus* cultivation (Shah et al., 2020), a potential growing disorder by continuous cropping (Santhanam et al., 2015). Second, according to the differences in root microbiomes among C treatments (Figure 3), irrigation using underground water containing salt in the F5S fields could change soil microbial communities in that field. The higher α-diversity of MIX and MIX locations intermediate from F5C to F5S in β- diversity could confirm the microbiome differences between the soils from F5C and F5S (Figures 2 and 3; Supplemental Figure S3). In addition, many microbes depended on both C and E effects (Supplemental Figure S4), suggesting that salt treatment changed the soil microbiome in F5S from the original microbiome, which did not differ between F5C and F5S. Therefore, the present study could reproduce the combination of the microbial community that changed with the environment and the environmental conditions that contributed to the change.

We found that the host genotype [G] significantly affected the α- and β-diversity of root microbiomes; nevertheless, the effect size was smaller than that of the C and E factors (Table 1). While previous studies have commonly shown a low contribution of host genotypes to shaping their root microbiome structures (Lundberg et al., 2012; Peiffer et al., 2013), assessing the impact compared to C and E is uncommon. In this study, even if a C- or E-dependent host effect existed, the sum of genotype-related effects on the root microbiome (17%) was smaller than the sole effect of the encountered community [C] (21%) and growth environment [E] (22%). This result indicates that many variations in root microbiomes could be defined by the microbial communities and growth environments encountered. In addition, 1776, 876, and 685 out of 3700 microbial strains were independent of the G-related (G, G × C, G × E, and G × C × E), C-related (C, G × C, C × E, and G × C × E), and E-related (E, G × E, C × E, and G × C × E) effects, respectively. These results indicate that many microbes that constitute the root microbiome were unaffected by host differences.

Furthermore, when the C and E were fixed, the 25-40% variation in microbial communities could be explained by G, and these effects were higher in salt conditions than in non-salt (Supplemental Figure S12A-F; Supplemental Table S5). This finding suggests that G effects could be growth conditions dependent. Therefore, small but significant host genotype effects on root microbiome suggested that the interaction with specific microbes could be controlled by the variable genetic basis of *Lotus* accessions depending on the growing conditions.

We observed a few microbial taxa, including *Pseudomonas*, *Sphingobium*, *Ralstonia*, and *Delftia*, which were sensitive to differences in plant genotypes. This finding is not surprising since the bacteria belonging to these genera have been reported as plant-interacting bacteria (Vishwakarma et al., 2020; Pfeiffer et al., 2017; Wozniak et al., 2019). The genus *Pseudomonas* is ubiquitous in diverse ecological habitats and encompasses plant symbionts and pathogens (Jain and Das, 2016). *Sphingobium* was highly abundant in the rhizosphere of maize and has been reported to show disease suppression ability for *Arabidopsis* (Innerebner et al., 2011). On the other hand, *Ralstonia* and *Delftia* bacteria were enriched as sensitive microbes to the interaction effects among G, C, and E. *Ralstonia* is a significant plant pathogen (Alvarez et al., 2019). In addition, several strains belonging to *Delftia* have been reported as plant growth-promoting rhizobacteria (Suchan et al., 2020). Bacteria under the direct influence of G factors were enriched in *Pseudomonas* and *Sphingobium*; nevertheless, their frequency was not correlated (Supplemental Figures S13 and S14). *Ralstonia* was observed to be more frequent in salty environments but less frequent in a genotype-dependent manner. *Delftia* strains were enriched in F5C inoculants, and their frequencies were higher, particularly MG20 and Gifu, under salt conditions. These results indicate that the affected genotypes differed among bacteria from the same genus, and their impact depended on the growth environment. Furthermore, these sensitive genera were not the predominant taxa in this study (*Pseudomonas*: 0.9%, *Sphingobium*: 2.5%, *Ralstonia*: 0.7%, and *Delftia*: 5.8%), supporting the low contribution to shaping microbiome structures. In this study, we found that genomic differences within plant species do not alter the structure of the root microbiome; nevertheless, there were bacteria whose interactions were under plant genotype-dependent control.

*Lotus*-microbe interactions depending on the plant genotypes could be more important for plant growth than the differences among encountered microbes (Table 2). The smaller effect of C on plant phenotypes, particularly plant shoot phenotypes, than that of G × C indicated that most altered microbes themselves had little effect on plant phenotypes; however, their interaction with G had more significant outcomes. These effects differed between plant shoot and root phenotypes; the root phenotype was more sensitive to differences in the encountered microbiome than the shoot phenotypes. This result suggests that plant roots interacted with and responded to soil microbiomes. In contrast, their interaction effects could be buffered/facilitated by each host genotype and spread into the shoots. Meanwhile, we could underestimate the impact of different microbial communities on plant phenotypes since inoculant microbe differentiation was limited due to their specific origin. Using natural habitats will have more diverse microbes and interactions between plant genotypes and microbes and help unravel the natural C and G × C effects on plant phenotypes.

There are two possible scenarios to explain why the G × C effect occurred: one is that the bacteria that affect plant phenotypes in a genotype-dependent manner are distributed differently in each inoculant community; another is that the genotype-dependent effects of bacteria on plant phenotypes are caused by each inoculant community, even if there are no differences in bacterial existence among inoculants. Even though around 70% of bacterial strains were distributed in different inoculant communities (Supplemental Figure S4) and could support the former scenario, it is still challenging to determine which scenario each G × C-related strain would follow. It was challenging to evaluate the effect of each bacterium on plant phenotypes because G, C, and E and their interactions affected both phenotypes and root microbiome and did not allow us to separate them. In this study, the root microbiome structure could explain 55% of the variation in plant shoot length, except for variance caused by the sole G factor. Accordingly, more detailed experiments and analyses, such as inoculation studies using synthetic communities (Finkel et al., 2020), will be more efficient in clarifying which microbes can affect plant phenotypes.

The genetic basis underlying the effects of G and G × C on interactions with microbes remains unclear. According to previous research, differences in plant genomes and the dissimilarity of their root microbiomes correlate with each other (Bouffaud et al., 2014; Schlaeppi et al., 2014; Terrazas et al., 2020). A higher correlation was observed at higher taxonomic levels (Wang and Sugiyama, 2020), and a lower correlation was observed for closely related species or within species levels (Terrazas et al., 2020). This finding suggests that the accumulated genomic divergence of plants may cause differentiation of the root microbiome. In contrast, there were no correlations between the kinship of *Lotus japonicus* and the root microbiome in this study (Supplemental Figure S2). By focusing on the natural diversity of *Lotus japonicus*, we can elucidate the genetic basis underlying the effects of G, G × C, and G × C × E on plant phenotypes and root microbiomes. This approach would be valuable for the disentanglement of the shape and maintenance of plant-microbiome interactions in nature.

## Materials and Methods

### Cross-inoculation experiments

We performed a cross-inoculation experiment to quantify the effects of plant genotype [G], soil microbiomes [C], growth environment [E], and their interactions on plant phenotypes and root microbiomes. We cultivated nine *Lotus* accessions with four soil microbial treatments (three microbes and a non-microbe control) under two conditions, resulting in 72 combinations.

We used eight *Lotus japonicus* natural accessions (Gifu, MG11, MG20, MG46, MG56, MG63, MG67, and MG68) and one *Lotus burttii* (B-303) for the cross-inoculation experiment. Three (Gifu, MG20, and *burttii*) were chosen because of their previous use as experimental lines (Kawaguchi et al., 2001; 2005). The other accessions were selected based on their genomic relationships and were referred to as Group 1 (MG67 and MG68), Group 2 (MG56 and MG63), and Group 3 (MG11 and MG46) (Shah et al., 2020). Seeds of *Lotus* accessions were obtained from the National BioResource Project in Japan.

Soil microbiomes were obtained from soils collected at the Kashimadai fields of Tohoku University (38.46 °N, 141.09 °E), located in northern Japan, in May 2020. Soil samples were obtained from two adjacent plots (F5C and F5S) where Lotus japonicus was cultivated for the last three years. Irrigation treatment was conducted using underground water containing salt at approximately 1/4 the concentration of seawater to F5S and regular water for F5C from 2017 to 2019. We separated 250 g of each soil and crushed it using a mixer with 250 mL of cold PBS buffer. Crushed soils were precipitated by centrifugation at 1000 × g for 10 min at 10 °C, and the supernatants were collected. The precipitates were returned to the mixer and crushing and centrifugation were repeated three times. The collected supernatant was filtered using Advantec 5A filter paper (collected particle size > 0.7 μm; ash content <0.01%). The filtered solutions were centrifuged at 8000 × g for 20 min at 10 °C, and the precipitates were collected. The precipitated product was diluted with 250 mL of PBS to obtain 1 mL/g microbial community extract. We used microbial community extracts from F5C, F5S, and a 1:1 mixture (MIX) for cross-inoculation experiments. As the difference in OD values between the extracts from F5C and F5S was less than 2%, we did not adjust their concentrations.

To set up the difference in the growth environment, we used two types of media for plant cultivation. Both were based on B&D medium (Broughton and Dilworth, 1971), and 1 mM KNO_3_ was added to the media to limit symbiosis with nitrogen-fixing nodule bacteria so that plant growth would not depend on them. NaCl was then added to the medium at a final concentration of 100 mM. The extracted microbial communities were added to each medium at a concentration of 1% (v/v), making eight different media (inoculants: F5C, F5S, MIX, and non-inoculant; media: SALT and non-SALT), which were used in the following experiments.

Partly scrubbed *Lotus* seeds were sterilized by immersion in 2% sodium hypochlorite for 3 min and rinsed three times with sterile MilliQ water. After overnight imbibition, the swollen seeds were sown on 1% agar plates, incubated in the dark for three days at 25 °C, and then grown at the same temperature under 16/8 light/dark conditions for 24 h. The rooted plants were transplanted into pots with a lid, filled with 300 mL sterilized vermiculite and 250 mL media, and grown at 25 °C under the same light conditions for four weeks. The growth pots were closed with lids to prevent cross-contamination. Two pots were used for each plant-inoculant-condition combination. A total of 144 pots (nine plant accessions × four inoculants × two conditions) were simultaneously grown in a growth chamber. We arranged the 144 pots into 10 groups of 14-15 pots each, and the locations of the groups were randomized weekly to prevent uneven lighting conditions. The group to which the pot belonged and the position of the pot in the group were randomly determined. Six plants were cultivated in each pot.

We then harvested whole plant bodies, imaged all individuals with a high-resolution scanner, and separated their roots and root nodules. Shoot length (SL), number of leaves (NOL), number of branches (NOB), and root length (RL) were measured from the scanned data as plant phenotypes. The roots were washed with sterilized distilled water, frozen in liquid nitrogen, and preserved at -80 °C until DNA extraction.

### DNA extraction for Miseq sequencing

Prior to DNA extraction, we cut each root sample into approximately 2 cm pieces and collected them randomly into sterilized tubes. The genomic DNA of each root sample was extracted using a Qiagen MagAttract 96 DNA Plant Core Kit (QIAGEN Inc., Valencia, CA, USA) according to the manufacturer’s instructions.

Pair-end library preparation for MiSeq sequencing was conducted using the two-step tailed PCR method described on Illumina (Illumina, San Diego, CA, USA). We used the following primer pairs to amplify partial sequences of the 16S rRNA gene:

V5F_MAUI_799 (forward): 5’-TCGTCGGCAGCGTCAGATGTGTATAAGAGACAGNNNHNNNWNNNHAACMGGATTAGATACCCKG-3’

V7R_1192 (reverse): 5’-GTCTCGTGGGCTCGGAGATGTGTATAAGAGACAGACGTCATCCCCACCTTCC -3’

The 3’ end to 18 bases and 19 bases of each primer (forward and reverse, respectively) were 16S rRNA universal sequences (Chelius and Triplett 2001). The 19-30 base from the 3’ end of the forward primer was the unique molecular identifier (Fields et al., 2020). The other regions were the Illumina overhang adapter sequences. The second-round PCR was performed using primer pairs with 16 unique indices: D501-D508 and A501-A508 (forward) and D701-D712 and A701-A712 (reverse) (Illumina).

The DNA concentration of the purified PCR products was adjusted and pooled into two different tubes, as the MiSeq run was performed in two separate runs. The samples used for the experiments are listed in Table S1. The paired-end libraries were mixed with 3% PhiX DNA spike-in control and used for sequencing on the MiSeq platform using Illumina MiSeq v.3 Reagent kit for 2 x 300 bp PE.

### Data analysis for microbiome

We conducted quality control for sequenced reads and paired-end read assembly using PEAR v0.9.6 (Zhang et al., 2014). The low-quality tails of each read were trimmed with a Phred score of 20 as the threshold, and trimmed reads with lengths of less than 200 bp were discarded. Then, pair-end reads with an overlap of more than 10 bp and a total length of more than 300 bp were combined. The UMI, primer, and target sequence regions of each read were identified based on the length of the sequences. The UMIs were counted without duplication, and the abundance of each sequence was determined based on the number of UMIs. Sequences with an average number of UMIs greater than or equal to 0.1 for all samples were chosen and used in the following analyses. The identification and counting of UMIs were performed using Python 3 in-house scripts.

A BLAST search (Camacho et al., 2009) was conducted for each sequence with the database containing the RDP11 bacterial 16S rRNA sequences (Cole et al., 2014) and *Lotus japonicus* Gifu genome v1.2 (Kamal et al., 2020) to assign the sequences to microbial taxa. A part of the classification rank of RDP11 that was out of alignment was manually corrected. All taxa were compared with the List of Prokaryotic names with Standing in Nomenclature (Parte et al., 2020) to prevent misclassification, and those that did not match were marked as “NotAssigned”. For the BLAST results, multiple sequences had the highest match rate, and the one with the exact genus name or the earliest RDP ID was selected. The bacterial community composition was reconstructed by excluding sequences whose top hits were the Gifu genome from the BLAST results.

### Microbial community analysis

To evaluate the effects of combinations of plant genotype [G], inoculant microbes [C], growth conditions [E], and their interactions with the plant root microbiome, we performed community analyses. The following analysis was performed using data aggregated from sequences with exact BLAST top hits.

Prior to analysis, to reduce biases due to differences in sampling depth, we subsampled community data based on the rarefaction curve using the *rarefy* function implemented in vegan, R (Dixon 2003). We identified the sample with the lowest slope at the endpoint of its rarefaction curve and adjusted the number of reads so that the slope at the endpoint of the rarefaction curve in all samples matched (Chao and Jost, 2012). We converted the rarefied community data into frequency data.

We then evaluated α-diversity of each sample using the Shannon diversity index (Shannon and Weaver, 1949) and calculated the effects of G, C, and E on diversity using a generalized linear model (GLM). In the GLM, α-diversity was the response variable, and the effects of G, C, and E and their interactions were explanatory variables. We chose the gamma distribution as the error distribution and log-link function for the model. Statistical significance was evaluated using the F-test. For the significant variables in the F-test (P < 0.05), we conducted the Tukey-Kramer test to compare α-diversities among all groups for that variable. We used the vegan packages (Oksanen, 2020) to calculate diversities and used the *glm*, *Anova*, and *glht* function in R (R core team, 2019) to estimate the effects of G, C, E, and interactions.

To distinguish the root microbiome structures, we calculated β-diversities among samples using the Morisita-Horn index (Horn, 1966). To visualize the similarity of microbial communities, we conducted non-metric multidimensional scaling (nMDS) analysis using the *metaMDS* function in vegan R (Oksanen et al., 2020) with 100 random parameters. We used PERMANOVA with the *adonis* function in vegan, R (Oksanen et al., 2020) with 99,999 permutations to evaluate which factors shape the bacterial community structure. To estimate the effects of variation within *L. japonicus* species on the root microbiome, we performed β-diversity analyses using the data, except for *L. burttii.* These analyses were also conducted for each inoculant-condition combination to clarify the effects of host genotype in different combinations.

Furthermore, we investigated the correlation between host genomes and root microbiome differences in each inoculant-condition combination using the Mantel test implemented in ape packages in R (Paradis and Schliep, 2019). The genetic distances of *Lotus japonicus* genomes were calculated using identical-by-state kinships based on the population genome information reported by Shah et al., (2020). The pairwise similarity distances of microbial communities were calculated using the Morisita-Horn index, which was calculated by averaging the microbial communities for each host.

We estimated the effects of G, C, and E and their interactions on the frequency of individual bacteria with GLM. We selected bacteria observed in more than six plant individuals for analysis to exclude excessive results from bacteria with minor distributions. In the GLM, each bacterial frequency was the response variable, and the effects of G, C, and E and their interactions were explanatory variables. We chose the gamma distribution as the error distribution and log-link function for the model. Statistical significance was evaluated using the F-test. Fisher’s exact test was used to evaluate whether the significantly affected strains were distributed disproportionately in specific genera and families. The GLM, F-test, and Fisher’s exact test were performed using R3.6.1.

### Data analysis for plant phenotypes

We first generated heatmaps using the host-standardized phenotypic values, whose mean values of each host genotype were set to zero to visualize the variation in phenotypes. We estimated correlations among phenotypes using Pearson’s product-moment correlation. To detect the effect of G, C, and E on the correlation among phenotypes, we performed a correlation test separately for each G, C, and E group. The heatmaps were illustrated by the *heatmap.2* program implemented in *gplots* in R3.6.1 (R Core Team, 2019). Correlation analyses were performed with the function implemented in ggpairs of R3.6.1 (R Core Team, 2019).

To analyze the effects of G, C, and E and their interactions on plant phenotypes, we used a generalized linear model (GLM). We used GLM instead of analysis of variance (ANOVA) because the distribution of phenotypic values deviated significantly from a normal distribution (Shapiro-Wilk test, P-value < 0.05 for all phenotypes). In the GLM, each phenotype was the response variable, and the effects of G, C, and E and their interactions were explanatory variables. We chose the gamma distribution as an error distribution and log link function for all phenotypes, because the distribution did not deviate from the expected distribution. We calculated the type II sums of squares for each variable, evaluated their statistical significance using F-tests, and estimated each variable’s effect size (η^2^). In addition, we performed the Tukey-Kramer test to compare plant phenotypes among the G, C, and E groups. These analyses of variance were performed with the *Anova* function implemented in the *car* library (Fox and Weisberg, 2019) and the *etaSquared* function implemented in the *lsr* library in R.3.6.1 (R Core Team 2019). The Tukey-Kramer test was performed with the *glht* function implemented in the multcomp library in R.3.6.1 (Hothorn et al., 2008). We performed the same analysis using a dataset that excluded non-inoculated individuals to evaluate the effect of differences in the inoculation community.

In addition, we performed the following statistical analyses to deal with the potential confounding factors caused by each pot because the individual plants in the same pot shared a unique environment. We evaluated the interclass correlation coefficients (ICC: variance between pots/all variance) of pots for each combination of inoculation tests (72 G × C × E combination) with the *glmer* function in R3.6.1. The ICCs were calculated with two plant phenotypes, plant shoot length, and root length, owing to low variance in the other phenotypes. Even if there was bias due to the combination of pot effects, the multi-level analysis containing pot information as a random effect was unsuitable because the pot variables completely masked the combination information. We randomly selected one of the pots from each combination to exclude pot bias, then evaluated G, C, and E, and their interaction effect using the GLM model for 1,000 permutations.

As both plant phenotypes and microbial communities depend on the effects of G, C, E, and their interactions, we calculated the extent to which root microbiome structures explained the variance in plant shoot length. We calculated the variance component using the following equation:

Y = u + e.

Y is an SL vector standardized for each host accession, and ε is an error term. μ is the similarity matrix of the root microbiome based on 1 - the Morisita-Horn similarity index matrix and the identical matrix used in the community analysis. We used the *emma* function in the R pipeline to calculate the variance component of u (Kang et al., 2008).

## Acknowledgments and Funding

Wild accessions of *L. japonicus* used in this research were provided by the National BioResource Project (“Lotus/Glycine”) of the Ministry of Education, Culture, Sports, Science, and Technology, Japan. This work was supported by JSPS KAKENHI [grant number 21K14763 to MB], JP20H2884 to SS, and the InRoot project coordinated by Jens Stougaard supported by The Novo Nordisk Foundation Grant Number NNF129SA0059362, Denmark.

## Author Contributions

Conceptualization, S.S and S.U.A.; Methodology, M.B. and S.S.; Investigation, M.B. and Y.A.; Analysis, M.B.; Data management, T.Y.A; Writing-Original draft, M.B. and S.S.; Writing-Review and Editing, S.S., S.U.A., and J. Q.; Supervision, S.S. and S.U.A; Funding Acquisition, M.B., S.S., and S.U.A.

## Supplemental Figure legends

**Supplemental Figure S1. Rarefaction curve.**

The rarefaction curve was derived from root microbiome data. The vertical axis represents the number of the strains. The horizontal axis represents the read count. Two clusters could be recognized from the sample size, which corresponded to the first and second runs of the MiSeq sequence.

**Supplemental Figure S2. Correlations between root microbiome structures and plant genetic distances.**

The vertical axis represents the root microbiome similarity based on the Morisita-Horn index. Prior to calculating diversity, the root microbiomes observed in each plant genotype with each inoculant-environment combination were averaged. The horizontal axis represents plant genetic similarity based on 1 – identical by state kinships among plant genotypes, based on Shah et al. (2020). Regression lines were drawn using ggplot with the *lm* function.

**Supplemental Figure S3. Effects of G, C, E, and their interactions on bacterial frequencies.**

The generalized linear model evaluated the effects of G, C, E, and their interactions on 3,700 bacterial strains, and the distributions of their significance (P values) are shown as violin plots. Dashed lines indicate P values equal to 0.05.

**Supplemental Figure S4. Venn diagram showing how many bacteria are affected by G, C, and E and how much the effects overlap.**

Venn diagrams of the significant effects of G, C, and E and their interactions on bacterial frequencies. Each cell number represents the number of bacterial strains affected. (A) Comparison of sole effects of G, C, and C, (B) comparison of G-related effects, and (C) comparison of C- and E-related effects.

**Supplemental Figure S5. Correlation and distribution of phenotypes in plant genotypes [G].**

Violin plots and histograms show the distribution of phenotypic values. The x- and y-axes in each scatterplot represent the following phenotypic values: shoot length, root length, number of leaves, and number of branches. Pearson’s correlation coefficients were calculated for all phenotypes (black indicates all groups).

**Supplemental Figure S6. Correlation and distribution of phenotypes in inoculants [C].**

Violin plots and histograms show the distribution of phenotypic values. The x- and y-axes in each scatterplot represent the following phenotypic values: shoot length, root length, number of leaves, and number of branches. Pearson’s correlation coefficients were calculated for all phenotypes (black indicates all groups).

**Supplemental Figure S7. Correlation and distribution of phenotypes in environments [E].**

Violin plots and histograms show the distribution of phenotypic values. The x- and y-axes in each scatterplot represent the following phenotypic values: shoot length, root length, number of leaves, and number of branches. Pearson’s correlation coefficients were calculated for all phenotypes (black indicates all groups).

**Supplemental Figure S8. Quantile-quantile plots for phenotypes of the cross-inoculation experiment to visualize the fits with the Gamma distribution.**

The shaded region represents 95% confidence intervals. (A) Shoot length, (B) root length, (C) number of leaves, and (D) number of branches.

**Supplemental Figure S9. Quantile-quantile plots for phenotypes of the cross-inoculation experiment, except for non-inoculant data, to visualize the fits with the Gamma distribution.**

The shaded region represents 95% confidence intervals. (A) Shoot length, (B) root length, (C) number of leaves, and (D) number of branches.

**Supplemental Figure S10. The significant effects of G, C, E, and their interactions on plant phenotypes in the randomized test to assess the pot effects in our cross-inoculation experiments.**

The vertical axis represents the log10 P-values for each effect. Dashed lines indicate P values of 0.05. (A) Shoot length, (B) root length, (C) number of leaves, and (D) number of branches.

**Supplemental Figure S11. Effects of G, C, E, and their interactions on plant phenotypes in the randomized test to assess the pot effects in our cross-inoculation experiments.**

The vertical axis represents the η^2^ values for each effect. (A) Shoot length, (B) root length, (C) number of leaves, and (D) number of branches.

**Supplemental Figure S12. Root microbiome structure based on β-diversity in each inoculant-condition combination.**

Non-metric multidimensional scaling (NMDS) for *Lotus* root microbiome dissimilarity (Morisita-Horn index) is shown. nMDS for the *Lotus* root microbiome for each inoculant-condition combination. The color represents different plant genotypes, and the areas of the identical genotypes are encompassed.

**Supplemental Figure S13. Sensitive genera to plant genotype and their correlation in the genus.**

The horizontal axes represent Spearman’s rank correlation coefficient, R, between the two bacterial strains significantly affected by plant genotype-related effects. The vertical axes represent the frequencies of the bacterial strain pairs. (A and B) *Pseudomonas* and *Sphingobium* were sensitive to the G effect. (C) *Ralstonia* is sensitive to G × C effects. (D) *Delftia*is sensitive to G × C × E.

**Supplemental Figure S14. Sensitive genera to plant genotype-related effects and their distributions in the root microbiome.**

The horizontal axes represent bacterial strains belonging to each genus. The vertical axes represent (A, B) plant genotype, (C) genotype × inoculant, and (D) genotype × inoculant × environmental combinations. Each cell color indicates the average bacterial frequency standardized for each bacterial strain.

## Reference

Alegria Terrazas R, Balbirnie-Cumming K, Morris J, Hedley PE, Russell J, Paterson E, Baggs EM, Fridman E, Bulgarelli D (2020) A footprint of plant eco-geographic adaptation on the composition of the barley rhizosphere bacterial microbiota. Sci Rep 10: 1–13

Álvarez B, López MM, Biosca EG (2019) Biocontrol of the Major Plant Pathogen *Ralstonia solanacearum* in Irrigation Water and Host Plants by Novel Waterborne Lytic Bacteriophages. Front Microbiol 10: 1–17

Bamba M, Aoki S, Kajita T, Setoguchi H, Watano Y, Sato S, Tsuchimatsu T (2019) Exploring genetic diversity and signatures of horizontal gene transfer in nodule bacteria associated with *Lotus japonicus* in natural environments. Mol Plant-Microbe Interact. doi: 10.1094/MPMI-02-19-0039-R

Bamba M, Aoki S, Kajita T, Setoguchi H, Watano Y, Sato S, Tsuchimatsu T (2020) Massive rhizobial genomic variation associated with partner quality in *Lotus*– *Mesorhizobium* symbiosis. FEMS Microbiol Ecol 1–15

Bamba M, Kawaguchi YW, Tsuchimatsu T (2018) Plant adaptation and speciation studied by population genomic approaches. Dev Growth, Differ 61: 12–24

Berendsen RL, Pieterse CMJ, Bakker PAHM (2012) The rhizosphere microbiome and plant health. Trends Plant Sci 17: 478–486

Bouffaud ML, Poirier MA, Muller D, Moënne-Loccoz Y (2014) Root microbiome relates to plant host evolution in maize and other Poaceae. Environ Microbiol 16: 2804–2814

Bouskill NJ, Lim HC, Borglin S, Salve R, Wood TE, Silver WL, Brodie EL (2013) Pre-exposure to drought increases the resistance of tropical forest soil bacterial communities to extended drought. ISME J 7: 384–394

Broughton WJ, Dilworth MJ (1971) Control of leghaemoglobin synthesis in snake beans. Biochem J 125: 1075–1080

Brown SP, Grillo MA, Podowski JC, Heath KD (2021) Correction to: Soil origin and plant genotype structure distinct microbiome compartments in the model legume *Medicago truncatula* (Microbiome, (2020), 8, 1, (139), 10.1186/s40168-020-00915-9). Microbiome 9: 1–17

Bulgarelli D, Rott M, Schlaeppi K, Ver Loren van Themaat E, Ahmadinejad N, Assenza F, Rauf P, Huettel B, Reinhardt R, Schmelzer E, et al (2012) Revealing structure and assembly cues for Arabidopsis root-inhabiting bacterial microbiota. Nature 488: 91–95

Busby PE, Peay KG, Newcombe G (2016) Common foliar fungi of *Populus trichocarpa* modify Melampsora rust disease severity. New Phytol 209: 1681– 1692

Camacho C, Coulouris G, Avagyan V, Ma N, Papadopoulos J, Bealer K, Madden TL (2009) BLAST+: Architecture and applications. BMC Bioinformatics 10: 1–9

Carrión VJ, Perez-Jaramillo J, Cordovez V, Tracanna V, De Hollander M, Ruiz-Buck D, Mendes LW, van Ijcken WFJ, Gomez-Exposito R, Elsayed SS, et al (2019) Pathogen-induced activation of disease-suppressive functions in the endophytic root microbiome. Science (80-) 366: 606–612

Chao A, Jost L (2012) Coverage-based rarefaction and extrapolation: Standardizing samples by completeness rather than size. Ecology 93: 2533–2547

Chelius MK, Triplett EW (2001) The diversity of archaea and bacteria in association with the roots of *Zea mays* L. Microb Ecol 41: 252–263

Cole JR, Wang Q, Fish JA, Chai B, McGarrell DM, Sun Y, Brown CT, Porras-Alfaro A, Kuske CR, Tiedje JM (2014) Ribosomal Database Project: Data and tools for high throughput rRNA analysis. Nucleic Acids Res 42: 633–642

de Vries FT, Williams A, Stringer F, Willcocks R, McEwing R, Langridge H, Straathof AL (2019) Changes in root-exudate-induced respiration reveal a novel mechanism through which drought affects ecosystem carbon cycling. New Phytol 224: 132–145

Enebe MC, Babalola OO (2020) Effects of inorganic and organic treatments on the microbial community of maize rhizosphere by a shotgun metagenomics approach. Ann Microbiol. doi: 10.1186/s13213-020-01591-8

Fields B, Moeskjær S, Friman VP, Andersen SU, Young JPW (2021) MAUI-seq: Metabarcoding using amplicons with unique molecular identifiers to improve error correction. Mol Ecol Resour 21: 703–720

Finkel OM, Castrillo G, Herrera Paredes S, Salas González I, Dangl JL (2017) Understanding and exploiting plant beneficial microbes. Curr Opin Plant Biol 38: 155–163

Fox J, Weisberg S (2019) An R Companion to Applied Regression, Third edit. Sage, Thousand Oaks

Gallart M, Adair KL, Love J, Meason DF, Clinton PW, Xue J, Turnbull MH (2018) Host Genotype and Nitrogen Form Shape the Root Microbiome of *Pinus radiata*. Microb Ecol 75: 419–433

Guo J, Ni BJ, Han X, Chen X, Bond P, Peng Y, Yuan Z (2017) Unraveling microbial structure and diversity of activated sludge in a full-scale simultaneous nitrogen and phosphorus removal plant using metagenomic sequencing. Enzyme Microb Technol 102: 16–25

Handberg K, Stougaard J (1992) *Lotus japonicus*, an autogamous, diploid legume species for classical and molecular genetics. Plant J 2: 487–496

Horn HS (1966) Measurement of "Overlap" in Comparative Ecological Studies. The University of Chicago Press for The American Society of Naturalists. http://www.jstor.com/stable/2459242 ECOLOGICAL STUDIES. 100: 419–424

Hothorn T, Bretz F, Westfall P (2008) Simultaneous inference in general parametric models. Biometrical J 50: 346–363

Innerebner G, Knief C, Vorholt JA (2011) Protection of *Arabidopsis thaliana* against leaf-pathogenic *Pseudomonas* syringae by *Sphingomonas* strains in a controlled model system. Appl Environ Microbiol 77: 3202–3210

Jain A, Das S (2016) Insight into the Interaction between Plants and Associated Fluorescent *Pseudomonas* spp. Int J Agron. doi: 10.1155/2016/4269010

Kamal N, Mun T, Reid D, Lin JS, Akyol TY, Sandal N, Asp T, Hirakawa H, Stougaard J, Mayer KFX, et al (2020) Insights into the evolution of symbiosis gene copy number and distribution from a chromosome-scale *Lotus japonicus* Gifu genome sequence. DNA Res 27: 1–10

Kawaguchi M (2000) *Lotus japonicus* “Miyakojima” MG-20: An early-flowering accession suitable for indoor handling. J Plant Res 113: 507–509

Kawaguchi M, Pedrosa-Harand A, Yano K, Hayashi M, Murooka Y, Saito K, Nagata T, Namai K, Nishida H, Shibata D, et al (2005) *Lotus burttii* takes a position of the third corner in the *Lotus* molecular genetics triangle. DNA Res 12: 69–77

Liu H, Brettell LE, Qiu Z, Singh BK (2020) Microbiome-Mediated Stress Resistance in Plants. Trends Plant Sci 25: 733–743

Lundberg DS, Lebeis SL, Paredes SH, Yourstone S, Gehring J, Malfatti S, Tremblay J, Engelbrektson A, Kunin V, Rio TG Del, et al (2012) Defining the core Arabidopsis thaliana root microbiome. Nature 488: 86–90

Mauchline TH, Malone JG (2017) Life in earth – the root microbiome to the rescue? Curr Opin Microbiol 37: 23–28

Naylor D, Degraaf S, Purdom E, Coleman-Derr D (2017) Drought and host selection influence bacterial community dynamics in the grass root microbiome. ISME J 11: 2691–2704

Oksanen J, Blanchet FG, Friendly M, Kindt R, Legendre P, McGlinn D, Minchin PR, O’Hara RB, Simpson GL, Solymos P, et al Vegan: Community Ecology Package. Version 2.5- 7. https://cran.r-project.org/web/packages/vegan/index.html

Paradis E, Schliep K (2019) Ape 5.0: An environment for modern phylogenetics and evolutionary analyses in R. Bioinformatics 35: 526–528

Parte AC, Carbasse JS, Meier-Kolthoff JP, Reimer LC, Göker M (2020) List of prokaryotic names with standing in nomenclature (LPSN) moves to the DSMZ. Int J Syst Evol Microbiol 70: 5607–5612

Peiffer JA, Spor A, Koren O, Jin Z, Tringe SG, Dangl JL, Buckler ES, Ley RE (2013) Diversity and heritability of the maize rhizosphere microbiome under field conditions. Proc Natl Acad Sci 110: 6548–6553

Pfeiffer S, Mitter B, Oswald A, Schloter-Hai B, Schloter M, Declerck S, Sessitsch A (2017) Rhizosphere microbiomes of potato cultivated in the high andes show stable and dynamic core microbiomes with different responses to plant development. FEMS Microbiol Ecol 93: 1–12

Rushworth CA, Song BH, Lee CR, Mitchell-Olds T (2011) Boechera, a model system for ecological genomics. Mol Ecol 20: 4843–4857

Santhanam R, Luu VT, Weinhold A, Goldberg J, Oh Y, Baldwin IT (2015) Native root-associated bacteria rescue a plant from a sudden-wilt disease that emerged during continuous cropping. Proc Natl Acad Sci U S A 112: E5013–E5120

Schlaeppi K, Dombrowski N, Oter RG, Ver Loren Van Themaat E, Schulze-Lefert P (2014) Quantitative divergence of the bacterial root microbiota in Arabidopsis thaliana relatives. Proc Natl Acad Sci U S A 111: 585–592

Shah N, Wakabayashi T, Kawamura Y, Skovbjerg CK, Wang M-Z, Mustamin Y, Isomura Y, Gupta V, Jin H, Mun T, et al (2020) Extreme genetic signatures of local adaptation during Lotus japonicus colonization. Nat Commun. doi: 10.1038/s41467-019-14213-y |

Shannon, C.E., Weaver W (1949) The mathematical theory of communication. Univ. Illinois Press 29:

Suchan DM, Bergsveinson J, Manzon L, Pierce A, Kryachko Y, Korber D, Tan Y, Tambalo DD, Khan NH, Whiting M, et al (2020) Transcriptomics reveal core activities of the plant growthpromoting bacterium delftia acidovorans RAY209 during interaction with canola and soybean roots. Microb Genomics 6: 1–13

Team RC (2019) R: A language and environment for statistical computing.

Trujillo ME (2016) Actinobacteria. eLS. doi: 10.1002/9780470015902.a0020366.pub2

Vishwakarma K, Kumar N, Shandilya C, Mohapatra S, Bhayana S, Varma A (2020) Revisiting Plant–Microbe Interactions and Microbial Consortia Application for Enhancing Sustainable Agriculture: A Review. Front Microbiol 11: 1–21

Wagner MR, Lundberg DS, Del Rio TG, Tringe SG, Dangl JL, Mitchell-Olds T (2016) Host genotype and age shape the leaf and root microbiomes of a wild perennial plant. Nat Commun 7: 1–15

Walters WA, Jin Z, Youngblut N, Wallace JG, Sutter J, Zhang W, González-Peña A, Peiffer J, Koren O, Shi Q, et al (2018) Large-scale replicated field study of maize rhizosphere identifies heritable microbes. Proc Natl Acad Sci 115: 7368– 7373

Wang B, Sugiyama S (2020) Phylogenetic signal of host plants in the bacterial and fungal root microbiomes of cultivated angiosperms. Plant J 104: 522–531

Weinert N, Piceno Y, Ding GC, Meincke R, Heuer H, Berg G, Schloter M, Andersen G, Smalla K (2011) PhyloChip hybridization uncovered an enormous bacterial diversity in the rhizosphere of different potato cultivars: Many common and few cultivar-dependent taxa. FEMS Microbiol Ecol 75: 497–506

Woźniak M, Gałązka A, Tyśkiewicz R, Jaroszuk-ściseł J (2019) Endophytic bacteria potentially promote plant growth by synthesizing different metabolites and their phenotypic/physiological profiles in the biolog gen iii microplate^TM^ test. Int J Mol Sci. doi: 10.3390/ijms20215283

Yadav AN, Verma P, Kumar S, Kumar V, Kumar M, Kumari Sugitha TC, Singh BP, Saxena AK, Dhaliwal HS (2018) Actinobacteria from Rhizosphere: Molecular Diversity, Distributions, and Potential Biotechnological Applications. New Futur Dev Microb Biotechnol Bioeng Actinobacteria Divers Biotechnol Appl 13–41

Yeoh YK, Paungfoo-Lonhienne C, Dennis PG, Robinson N, Ragan MA, Schmidt S, Hugenholtz P (2016) The core root microbiome of sugarcanes cultivated under varying nitrogen fertilizer application. Environ Microbiol 18: 1338–1351

Zhang J, Kobert K, Flouri T, Stamatakis A (2014) PEAR: A fast and accurate Illumina Paired-End reAd mergeR. Bioinformatics 30: 614–620

Zhang J, Liu YX, Zhang N, Hu B, Jin T, Xu H, Qin Y, Yan P, Zhang X, Guo X, et al (2019) NRT1.1B is associated with root microbiota composition and nitrogen use in field-grown rice. Nat Biotechnol 37: 676–684

